# Near equilibrium unbinding of streptavidin/biotin using single molecule acoustic force spectroscopy

**DOI:** 10.1101/2025.07.18.665553

**Authors:** Yogesh Saravanan, Lorenzo Villanueva, Christian Leveque, Mauro Modesti, Oussama El Far, Claire Valotteau, Felix Rico

## Abstract

The dissociation of the streptavidin-biotin (SA-b) bond has been widely characterized using bulk and single molecule force spectroscopy (SMFS) techniques. However, the dissociation rates (*k*_off_) from SMFS (∼10^−1^ s^-1^) typically do not align with those from bulk approaches (∼10^−6^ - 10^−5^ s^-1^), likely because SMFS measurements are conducted far from equilibrium. Near equilibrium SMFS requires high throughput measurements to obtain large enough statistics, and high stability over long time measurements at ultraslow loading force rates, impractical in most SMFS techniques. Here, we developed in-situ force calibration strategies for acoustic force spectroscopy (AFS) to probe the unbinding forces of SA-b in the near equilibrium regime, from 10 pN/s down to 10^− 3^ pN/s. The resulting *k*_off_ was in excellent agreement with that from bulk measurements on the very same system. Combined with our previous SA-b data, we covered 15 orders of magnitude in loading rate, setting the ground for SMFS over the widest dynamic range, essential to fully describe the energy landscape of biomolecular processes.

## MAIN TEXT

Receptor-ligand interactions govern essential biological processes, from cellular adhesion to antigen recognition^1^. Quantifying the unbinding kinetics of these interactions is thus important for understanding biological functions. While traditional bulk equilibrium methods have been widely used to estimate unbinding rates through concentration-based assays^2^, they offer limited insight into the molecular mechanics of dissociation. Single-molecule force spectroscopy (SMFS) has established as a powerful tool to probe the unbinding forces of receptor-ligand complexes, providing access to the energy landscapes that govern these interactions under force. Two primary SMFS modes — force-clamp and dynamic force spectroscopy (DFS) — allow probing unbinding kinetics by measuring, respectively, either unbinding times at constant force or unbinding forces at constant loading rate.

The bond formed by the tetrameric protein (strept)avidin (SA) and the small molecule biotin (b) is well-studied and widely used in molecular biology to tag or immobilize biomolecules. Ensemble measurements near equilibrium have revealed its high affinity (*K*_D_ ∼ 0.1 pM^-1^) and long lifetime (*k*_off_ ∼ 10^−5^ s^-1^) ^2–4^. The SA-b complex has been also widely probed using SMFS, first using atomic force microscopy (AFM), and later with other experimental techniques and molecular dynamics simulations^5–7^. This led to the establishment of theoretical frameworks based on the transition state theory to extract the parameters of the energy landscape: the distance to the transition state (*x*_β_) and, through extrapolation, the dissociation rate at zero force (*k*_off_) (Table S1)^7–11^. In this context, estimation of *k*_off_ using either ensemble or single molecule approaches should in principle lead to similar values. Nevertheless, despite the large number of SMFS works on the SA-b system, the *k*_off_ values have been systematically orders of magnitude apart from those using ensemble measurements^9,12–15^. This discrepancy may be explained by various factors, including the presence of multiple barriers across the energy landscape^13^, favored unbinding pathways at equilibrium^14,15^, and even maturation of the bond^16^.

To resolve this discrepancy, it is necessary to carry out SMFS experiments on the SA-b bond near equilibrium conditions, *i*.*e*. at ultraslow loading rates. However, given the extremely slow unbinding kinetics and the stochastic nature of SMFS measurements, conventional techniques such AFM are impractical due to their low throughput and force stability, making it difficult to obtain statistically relevant datasets. Acoustic force spectroscopy (AFS) is a high-throughput SMFS technique that uses standing acoustic waves to pull simultaneously on multiple beads anchored through the individual receptor-ligand complex of interest^17–19^. AFS allows force to be applied to several complexes in parallel, providing a large number of unbinding events in a single experiment. AFS has also shown good force stability over long times, thus practically allowing measurements at ultraslow loading rates. However, the acoustic field within the AFS chamber is not homogeneous, and careful force calibration is necessary for accurate measurements^20^. We have recently implemented an approach to calibrate each bead in real time during force-clamp measurements on individual bonds^21^. However, conventional force-clamp calibration appears impractical in experiments at constant loading rate, as it requires a pre-calibration step during which several beads are lost. Therefore, a new calibration approach. seems necessary to allow DFS measurements at ultra-slow loading rates.

In this study, we developed a method for accurately calibrating each bead during DFS measurements. This allowed us to pull on SA-b bonds over several hours at ultraslow, near equilibrium loading rates (∼10^−3^ pN/s) and up to conventional rates (∼10 pN/s). This near equilibrium regime led to *k*_off_ values in excellent agreement with those from bulk measurements. Our approach allowed us to extend the current loading rate range of the dynamic force spectrum by 4. Remarkably, combined with previous AFM measurements and molecular dynamics simulations we reached an overall range covering 15 orders of magnitude in loading rate.

We used a commercial AFS system mounted on the stage of a home built optical microscope with a 40X objective. The surface of the AFS chamber was coated with streptavidin by physisorption and silica beads covalently functionalized with DNA strands of 1765 base pairs (∼590 nm contour length) featuring a biotin at the end were injected in PBS buffer. After allowing the biotinylated DNA strands to interact with the SA on the surface, we modulated the acoustic power to apply a ramp of pulling force to the beads at a defined rate (*dP/dt* in units of % per second), which determines the loading rate applied to the bond (Fig 1 A-B). The recorded traces were processed and analysed using in-house Python-based software (https://github.com/DyNaMo-INSERM/PyAFSpectra).

**Figure 1:**
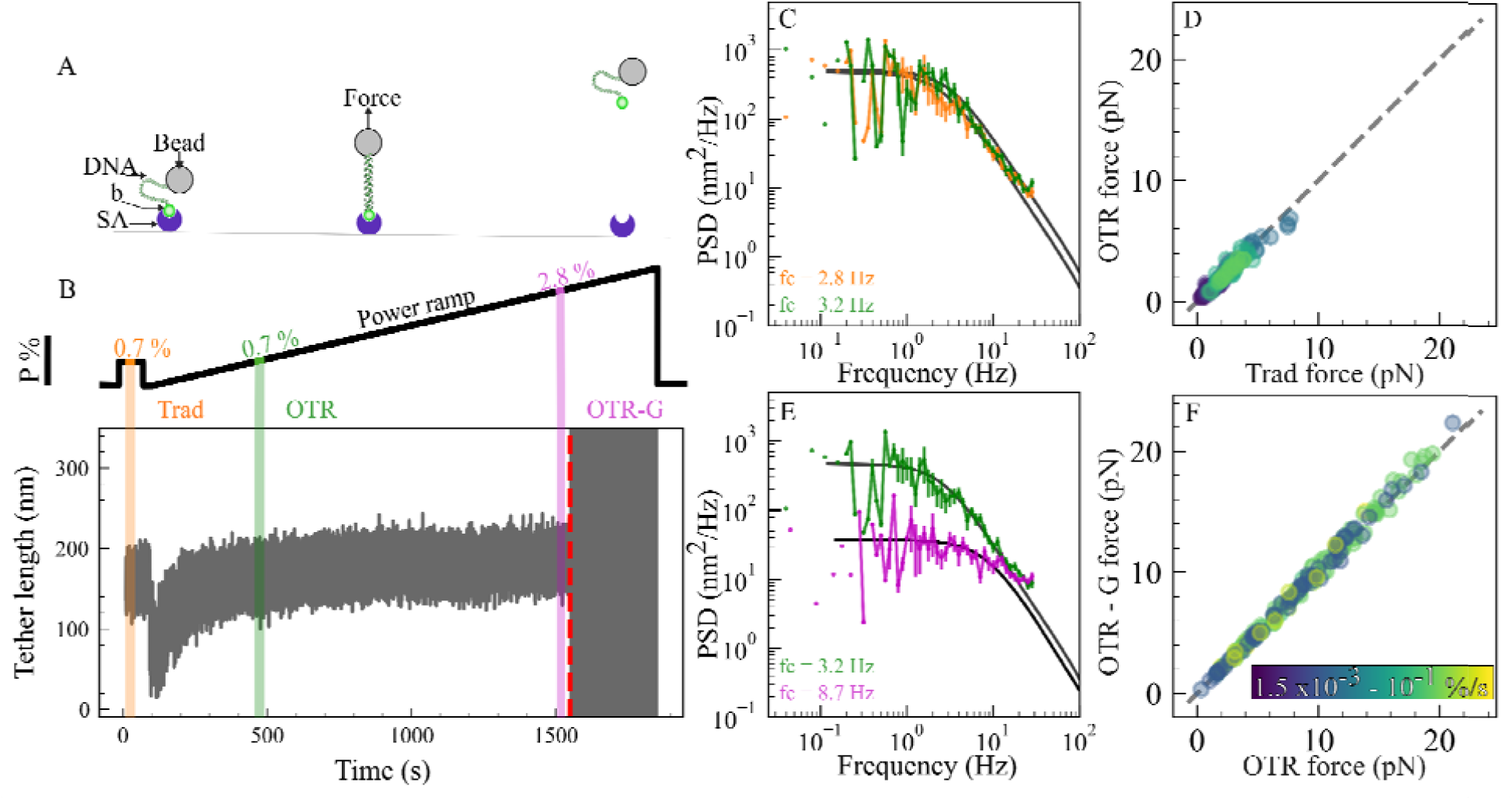
AFS pulling experiment and force calibration strategies. (A) Silica beads tethered through biotinylated (b) DNA interact with streptavidin (SA) physiosorbed on the surface of the AFS chamber (not to scale). When a ramp of acoustic force is applied, the beads, and therefore the streptavidin-biotin (SA-b) bonds, are pulled until the bonds rupture and the beads detach. (B) The bead movements are tracked in the X, Y, and Z directions, allowing us to calculate the evolution of the tether length, which increases when a power is applied. Once the bond has ruptured (red dotted line), the tracking signal is lost. To accurately estimate the bond rupture force and loading rate, force calibration is performed as close to rupture as possible. Power spectral densities (PSDs) are built along the force ramp from both the X and Y tracked data. Traditionally, the PSD is built over a constant acoustic power step (Trad, 0.7% of power), as depicted in orange. The new force calibration strategy on the ramp (OTR) relies on building a PSD over a short section of linear power ramp (in green) to obtain the bead diffusion coefficient (*D*) and trap stiffness (*k*). The OTR global strategy (OTR-G) relies on using the bead diffusion coefficient determined at low power and fitting only the trap stiffness at high power. (C) Comparison of PSDs built along the X-axis at constant power (traditional calibration, orange, *k* =(1.0□±□0.05)□×10^−3^ pN/nm, *D* =(6.50 ± 0.1)□×□10□□μm^2^/s, force = 1.8□±□0.1□pN) vs. OTR method (green, *k* = (0.80 ±□0.03)□×10^−3^ pN/nm, *D* = (10.8 ± 1.0)□×□10□□μm^2^/s, force = 1.4□±□0.2□pN) for a single bead at an equivalent average power (0.7%). (D) Comparison of the forces calibrated using the traditional (Trad) and OTR strategies along X and Y axes for 22 individual beads. The color scale indicates the power rate of the linear power ramp. The dotted line indicates the identity line for reference. (E) Comparison of PSDs built on the ramp along the X-axis at low power (0.7%, green) and higher power (2.8%, purple), showing the overlap of the diffusive tails, allowing to use an average diffusion constant (*D*_G_ = 5.18□×□10^4^□μm^2^/s) from low-power PSD fits to calculate the stiffness (*k* = (4.0□±□0.1)□×10^−3^ pN/nm) and force (.3□±□0.2□pN) at high power, even if the PSD corner frequency is closed to the limit. (F) Comparison of the force calibrated using the OTR and OTR-G strategies along X and Y axes for 7 individual beads tracked at high frame rate (120 Hz). The color scale indicates the power rate of the linear power ramp. The dotted line indicates the identity line for reference.

The forces experienced by the tethered beads were accurately calibrated for each bead. Conventional calibration involves acquisition of the thermal fluctuations of the bead in the XY plane at a constant power, constructing the power spectral density (PSD), and fitting a Lorentzian to determine the stiffness (*k*) and diffusion constant (*D*) (Eq S1). This calibration is generally performed prior to the experiment, through a dedicated step, during which the thermal fluctuations of the bead are acquired at different constant power levels, each applied for tens of seconds. The resulting PSDs allow the acoustic force to be determined for each power, generally through a constant force calibration factor per AFS chamber^17^. However, the acoustic force field within the AFS chamber is spatially heterogeneous and a single calibration value is not valid for all beads^20^. This makes force calibration a difficult although critical step in AFS. In addition, a significant proportion of single tethered beads (20-50%) are likely to detach during the initial calibration phase, effectively reducing the number of measured unbinding events. To overcome these limitations and still accurately estimate forces and loading rates, we propose an *in-situ* calibration strategy, “on the ramp” (OTR), which consists of bead calibration through fluctuations during the force ramp applied to the bonds. For that, we defined a 20 s time window (1200 points at a typical 60 Hz sampling rate) as close as possible to the unbinding event. A PSD of the position fluctuations was constructed and a Lorentzian was fitted to it to extract the stiffness *k* and diffusion constant *D* (Fig S1). Knowing the tether length *L* and bead radius *R* allowed us to accurately determine the average force over the time window, <*F*^OTR^> = *k (L+R)*, at the corresponding average applied power, <*P*^OTR^>. The unbinding force (*F*_u_) and its corresponding loading rate (*r*_*F*_), were deduced from the OTR calibrated force and the power at which the bond ruptured (*P*_*r*_) as *F*_u_ = <*F*^OTR^>/<*P*^OTR^>*P*_r_ and *r*_f_ = <*F*^OTR^>/<*P*^OTR^>d*P*/dt.

To assess the accuracy of the OTR calibration, we compared PSDs built using OTR with traditional PSDs built over constant force steps at equivalent powers (Fig 1C). The PSDs overlapped and led to similar values of *k* and *D*. The resulting forces from the OTR and traditional calibration strategies were in excellent agreement for several individual beads across the AFS chamber, over a wide range of power rates, from 1.5 × 10^−3^ to 0.1 %/s, equivalent to loading rates of 10^−3^ to 10^−1^ pN/s (Fig 1E). At the lowest rates, this represents pulling on individual bonds for over up to 3 hours prior to rupture, conditions we considered to be near equilibrium.

Despite its accuracy, the OTR strategy has limited application at the highest load rates. Indeed, to ensure a robust non-linear fit of both *k* and *D*, at least two-thirds of the PSD must describe the diffusive regime. Moreover, the Nyquist criterion states that the maximum frequency of the PSD should be half the camera frame rate^22^. This restricts the OTR fitting strategy to a maximum corner frequency (f_c_) of ∼10 Hz for a 60 Hz acquisition rate, and thus to relatively low forces at same experimental conditions (bead size, DNA length, camera frame rate). One way to overcome this limitation would be to increase the frame rate of our camera, but this would reduce the field of view and the number of tracked beads, and hence the experimental throughput. In order to extend the range of calibrated forces using the OTR strategy, we adapted to linear power ramp experiments the global PSD fitting method that we recently developed for force-clamp measurements^21^. This resulted in the OTR global strategy (OTR-G), consisting in determining an average *D*_*G*_ from PSDs built from windows at low forces (the start of the power ramp, green window on Fig 1), that satisfy the Nyquist criteria. This *D*_*G*_ value is then used to fit only *k* from the PSD built from windows at higher forces (pink window on Fig 1). To assess the accuracy of this calibration strategy, we acquired traces at high sampling rates (120 Hz), which allowed us to apply both OTR and OTR-G approaches. The forces obtained with both strategies showed excellent agreement (Fig 1E). Therefore, using the OTR-G calibration allows overcoming the Nyquist criteria at high forces while reducing the number of fitting parameters of the Lorentzian fit (Fig S2). This extended the range of measurable forces (10 pN to ∼50 pN) and loading rates (from 0.01 to 0.7 pN/s) while maintaining the same experimental conditions. To further increase this range, we recorded videos at 200 Hz, using a camera with faster frame rate. This gave us a sufficient sampling rate to apply the OTR-G strategy to traces with faster loading rates (up to 10 pN/s). The use of higher acoustic power revealed, however, a non-linear force versus power response (Fig S4). Earlier works have reported this effect, which may be attributed to a power-dependent shift in the resonance frequency^17,20^. To accurately determine the unbinding force and loading rate, we determined the forces using the OTR-G strategy across the full power ramp and then use the last linear part of the ramp (about one half) to determine the loading rate and to extrapolate the rupture force (Fig S5). This strategy enabled us to accurately determine the unbinding forces and loading rates up to ∼10 pN/s, reaching the lower limit of other experimental techniques such as AFM and allowing data comparison and combination to cover a wider dynamic range.

Using the OTR & OTR-G calibration strategies we accurately determined the unbinding forces of SA-b over four orders of magnitude in loading rate, from 10^−3^ pN/s to 10 pN/s, obtaining the near-equilibrium dynamic force spectrum (DFS) of this bond (Fig 2A, Fig S7). Most probable unbinding forces increased linearly with the logarithm of the loading rate over most of the spectrum, while they revealed a slight upturn at the highest loading rates. We applied the Bell-Evans model, which allows the determination of the energy landscape parameters. Since it assumes a linear dependence of the force to the logarithm of the loading rate, we fitted it to four consecutive linear fragments of the DFS^23^. The results suggested either a progressive shortening of the distance to the transition state (*x*_β_) or the presence of multiple barriers, ranging from ∼4 nm at the slowest loading rates to ∼1 nm at the highest (Fig 2 B), as well as an increase in the dissociation rate at zero force *k*_off_, rising from ∼10^−5^ s^-1^ to 10^−2^ s^-1^ (Fig 2C). Our previous work, which combined high-speed force spectroscopy (HS-FS) and molecular dynamics (MD) simulations, revealed two barriers and indirectly suggested the presence of a third barrier at a distance larger than ∼0.8 nm^13^, which would be in agreement with the outermost barrier at 4 nm. Importantly, the extrapolated dissociation rate at zero force from the slowest loading rate window (10^−4.2±1.5^ s^-1^), which is in good agreement with previous bulk measurements ^2–4,24^. To determine the bulk *k*_off_ in our very molecular system, we carried out surface plasmon resonance (SPR) experiments with the same molecules, immobilizing SA on the chip, capturing the soluble biotinylated DNA and monitoring dissociation kinetics The obtained *k*_off_ = (1.6 ± 0.9) x 10^−5^ s^-1,^ in good agreement with our single molecule AFS results (Table S2). This suggests that, unlike earlier experiments at higher loading rates, the loading rate regime explored by AFS probes the outermost barrier of the SA-b energy landscape, equivalent to the equilibrium unbinding pathway.

**Figure 2:**
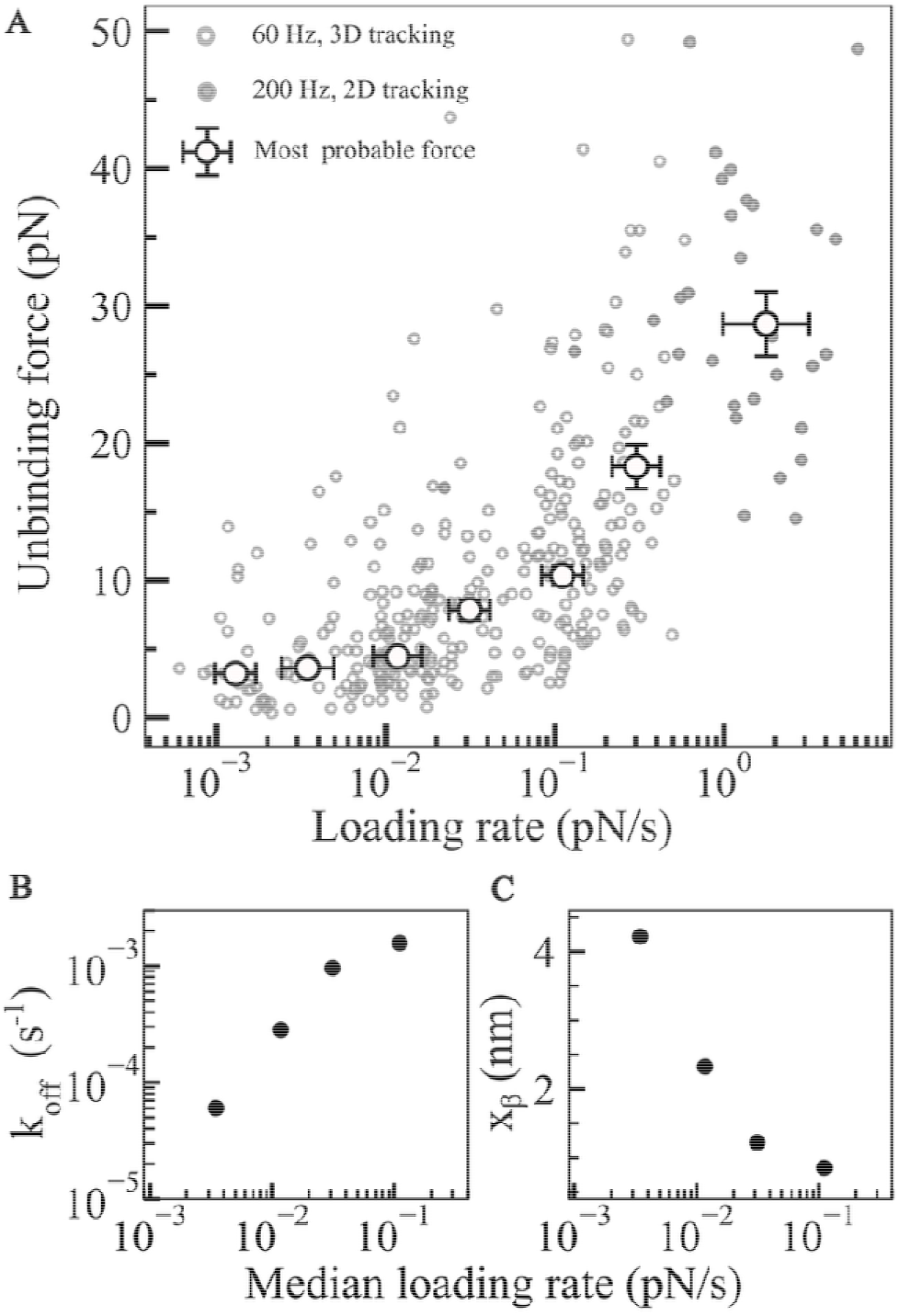
Near equilibrium dynamic force spectrum (DFS) of SA-b Interaction. (A) Rupture forces of the SA–b bond measured using AFS across 4 orders of magnitude in loading rate, at 60□Hz (open gray circles) and 200□Hz (closed gray circles). Most probable unbinding force (black circles) were obtained from Gaussian fits over discrete loading rate intervals; the geometric mean of the loading rate within each interval is shown, with error bars representing the standard deviation of the mean (Fig S6). Data were fitted using the Bell–Evans model over rolling windows of four consecutive points to extract the dissociation rate at zero force *k*_off_ (B) and the distance to the transition state *x*_β_ (C), plotted as a function of the median loading rate within each fitting window.

A recent SMFS work ensuring specific pulling geometry on the SA molecule found a dissociation rate of 10^−6^ s^−1^, similar to that obtained using ensemble measurements^12^. These results suggested that different unbinding pathways were explored if the pulling geometry was not defined, as in our AFS work, which would favor pathways with lower unbinding forces^15^. Our results confirm the idea that near equilibrium forced unbinding is dominated by the outermost barrier, likely the one dominating equilibrium unbinding.

The DFS we obtained revealed a certain curvature at the highest loading rates (Fig 2) which called for the application of alternative models^25–29^ (Fig S7, Table S3). One obvious approach was to consider the Friddle-Noy-De Yoro (FNdY) model, which was developed to capture both the near-equilibrium regime, and the activated regime ^28^. Even if this model fitted best our data (Fig S7), it is difficult to justify for receptor-ligand systems probed by AFS and provided unrealistic results (Table S3). First, the FNdY model assumes a constant stiff probe, which does not apply to AFS where the stiffness *k* of the acoustic trap is extremely soft and scales with the applied force *F*, gradually increasing from 10^−3^ to 10^−2^ pN/nm. Secondly, the model predicts a plateau at low loading rates, which was not clearly observed in our results. Thirdly, when the bond breaks, SA is physically separated from biotin, which makes the formation of a virtual trap unrealistic. Fourthly, the resulting parameters predict an extremely large free energy difference between the bound and unbound states of 500-5000 *k*_B_*T*, a distance to the transition state of 0.54 nm, and a *k*_off_ of 0.003 s^-1^ not in agreement with bulk measurements. Another recent theoretical study also considered reversible bond kinetics through the analysis of unbinding and rebinding events^29^. However, bead tracking in AFS lacks the temporal resolution and Z-tracking precision required and, thus, the approach was not applicable to our data. For these reasons, rebinding models were unsuitable for the SA-b complex and other approaches were evaluated.

Two other models predict curvature of the DFS by assuming that the applied force shifts the position of the transition state: the Cossio-Hummer-Szabo (CHS) and the Dudko-Hummer-Szabo (DHS) models. In the CHS model, the shortening of *x*_β_ is modulated by the kinetic brittleness parameter (μ). With a fixed brittleness of 0.01, the CHS model predicted a *k*_off_ of ∼10^−6^ s^-1^, in good agreement with our bulk SPR measurements. However, it predicted an unrealistic distance to the transition state of 4.8 nm, close the total length of one SA monomer. With a *k*_off_ fixed to the bulk 1.6 × 10^−5^ s^-1^ and free brittleness parameter, the CHS model fitted the DFS well and predicted a kinetic brittleness of 0.39 and a distance to the transition state of ∼3.0 nm, which is approximately the distance from the SA binding pocket to the outermost residue of loop 7-8 along the unbinding pathway. The earlier DHS model with a shape factor of the energy landscape ν=2/3, predicted a *k*_off_ of ∼2 × 10^−4^ s^-1^, in reasonable agreement with the bulk results, and a reasonable *x*_β_ of ∼2.1 nm, but the model failed to describe the spectrum at the highest loading rates. Interestingly, when the shape factor ν was left free, the DHS model fitted the spectrum better and predicted a reasonable *k*_off_ of 2.5×10^−6^ s^-1^ although the *x*_β_ of ∼5 nm was again unrealistically large, and ν fall in the disallowed range at ∼0.01. This low value of ν (ν < 0.5) was suggested by Hyeon to be the sign of the existence of multiple barriers across the unbinding pathway^23^, which justifies our first approach of piecewise fitting of the Bell-Evans model.

To obtain a complete picture of the full energy landscape, we combined our AFS results with previous data from HS-FS and MD simulations complemented with newly acquired AFM data at low loading rates (Fig 3, Fig. S8)^13^. The resulting DFS covers 15 decades in loading rate, from 10^−3^ to 10^13^ pN/s, making the SA-b bond the system explored over the widest dynamic range to date. Rupture forces reveal a clear curvature along the whole spectrum, which could not be adequately described by a single barrier. Based on the new AFS regime, we propose an energy landscape consisting of three barriers across the unbinding pathway at around 0.2, 0.5 and 4 nm.

**Figure 3:**
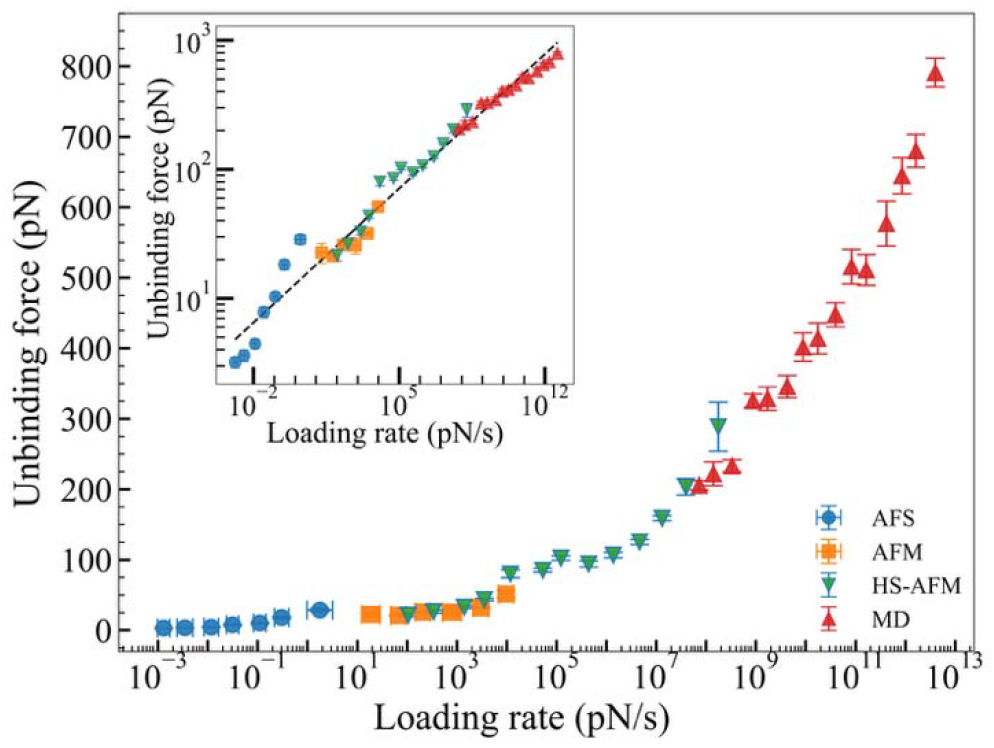
15 decades wide dynamic force spectrum of the SA-b interaction. Dynamic force spectrum (DFS) over 15 decades of loading rate using AFS (blue circles) combined with data from conventional (orange squares) and high-speed (green inverted triangles) atomic force microscopy^13^, and molecular dynamics simulations (red triangles). **Inset**: Log-log DFS of SA-b shows a weak power-law scaling (black dotted line) of the unbinding forces.

Therefore, as initially suggested by Evans and Ritchie in their seminal work, our results reinforce the idea that a complete description of the energy landscape requires exploration of the widest possible dynamic range. The long distance to the outermost barrier suggests a possible involvement of the 7-8 loop of the biotin-bound SA monomer or interaction with a neighbor SA monomer of the tetramer.

A log-log representation better reflects the overall response of the unbinding forces over loading rate. Interestingly, in this form, the DFS appears rather linear and is well described by a power law with an exponent of ∼0.14 (Fig 3 inset). Power laws commonly emerge from the existence of rough energy landscapes with multiple barriers of various heights leading to a continuous spectrum of relaxation rates. This may set a novel ground for further theoretical development.

In this work, we developed an approach based on AFS to probe rupture forces of individual bond at near equilibrium conditions. We introduced two *in situ* force calibration strategies, OTR and OTR-G, to accurately determine the rupture forces applied to each individual bead, regardless of the power applied or its spatial position within the chamber, without any assumption about a uniform (valid to all beads) and linear scaling relationship between the acoustic force and power. The strategies are applicable to other techniques, such as magnetic tweezers and enabled us to measure a wide range of forces covering a wide range of loading rates without changing the experimental conditions. This allowed us to probe the unbinding of the well-known SA-b complex. Our results show a DFS with clear curvature requiring multiple free energy barriers for a complete description. Importantly, the extrapolated *k*_off_ from the AFS is in excellent agreement with equilibrium bulk SPR measurement, solving a long-lasting disagreement between experimental techniques. Combined with previous and extended AFM-based experiments and MD simulations, we cover 15 orders of magnitude in loading rate and build an accurate description of the “reaction coordinate” at near and far from equilibrium conditions, suggesting the presence of at least three energy barriers along the unbinding pathway. Despite recent theoretical developments unifying near equilibrium, activated and deterministic loading rate regimes, available models with a single barrier do not fully describe the complete SA-b spectrum. Nonetheless, we believe that our results combined with previous data represents an invaluable dataset that will hopefully inspire the development of new theoretical frameworks. Our approach gives access to an unexplored dynamic regime that, together with other methodologies, will provide a more representative picture of the unbinding energy landscape of receptor-ligand complexes.

## ASSOCIATED CONTENT

**Supporting Information**. Materials and Methods.

Figures S1 to S8, Tables S1 and S2.

## AUTHOR INFORMATION

## Corresponding Authors

*

## Author Contributions

YS performed the AFS measurements and analyses under the supervision of CV and FR. LV contributed to code development. MM provided the DNA tethers. CL and OEL performed the SPR measurements. The manuscript was written through contributions of all authors. All authors have given approval to the final version of the manuscript.

## Funding Sources

The authors would like to acknowledge all the funding sources that supported this work.

This project received funding from the European Research Council (ERC) under the European Union’s Horizon 2020 research and innovation programme (grant agreement number 772257 to FR) and the Marie Curie Sklodowska action (MSCA-IF) (grant agreement no. 895819 to CV), from ITMO Cancer of Aviesan on funds Cancer 2021 (ATIP Avenir to CV) and from the Agence Nationale de la Recherche (ANR AAPG2023 - PRC – XXL to MM and FR). The AFS setup was acquired thanks to the grant Projet exploratoire region PACA 2017 – AcouLeuco cofunded by the ANR (ANR-15-CE11-0007-01 to FR). We also acknowledge Inserm, CNRS, and amU for regular support.

## ACKNOWLEDGMENT

We thank Alexandre Ortega, for fruitful discussions and his experimental help.

## ABBREVIATIONS

SA: streptavidin
b: biotin
AFS: acoustic force spectroscopy
AU: Arbitrary units
D: diffusion constant
DFS: dynamic force spectrum
DNA: deoxyribo-nucleic acid
f_c_: corner frequency,
f_s,_: camera sampling frequency
k,: spring constant
k_B_,: Boltzmann constant
T,: temperature
OTR,: on the ramp
OTR-G,: on the ramp global
PSD,: power spectral density
SMFS,: single molecule force spectroscopy.

## Notes

https://github.com/DyNaMo-INSERM/PyAFSpectra

### Competing Interest Statement

The authors have declared no competing interest.

## REFERENCES

[1] Alberts, B.; Johnson, A.; Lewis, J.; Raff, M.; Roberts, K.; Walter, P. Molecular Biology of the Cell, 4th ed.; Garland Science, 2002.

[2] Koussa, M. A.; Halvorsen, K.; Ward, A.; Wong, W. P. Nature Methods, 2, (2015), 123– 126.

[3] Hyre, D. E. Protein Science, 3, (2006), 459–467.

[4] Chivers, C. E.; Koner, A. L.; Lowe, E. D.; Howarth, M. Biochemical Journal, Pt 1, (2011), 55–63.

[5] Florin, E.-L.; Moy, V. T.; Gaub, H. E. Science, 5157, (1994), 415–417.

[6] Izrailev, S.; Stepaniants, S.; Balsera, M.; Oono, Y.; Schulten, K. Biophysical Journal, 4, (1997), 1568–1581.

[7] Merkel, R.; Nassoy, P.; Leung, A.; Ritchie, K.; Evans, E. Nature, 6714, (1999), 50–53.

[8] Bell, G. I. Science (New York, N.Y.), 4342, (1978), 618–627.

[9] Evans, E.; Ritchie, K. Biophysical Journal, 4, (1997), 1541–1555.

[10] Rico, F.; Moy, V. T. Journal of molecular recognition□: JMR, 6, (2007), 495–501.

[11] Guo, S.; Ray, C.; Kirkpatrick, A.; Lad, N.; Akhremitchev, B. B. Biophysical Journal, 8, (2008), 3964–3976.

[12] Johnson, K. C.; Thomas, W. E. Biophysical Journal, 9, (2018), 2032–2039.

[13] Rico, F.; Russek, A.; González, L.; Grubmüller, H.; Scheuring, S. Proceedings of the National Academy of Sciences, 14, (2019), 6594–6601.

[14] Sedlak, S. M.; Bauer, M. S.; Kluger, C.; Schendel, L. C.; Milles, L. F.; Pippig, D. A.; Gaub, H. E. PLOS ONE, 12, (2017), e0188722.

[15] Sedlak, S. M.; Schendel, L. C.; Melo, M. C. R.; Pippig, D. A.; Luthey-Schulten, Z.; Gaub, H. E.; Bernardi, R. C. Nano Letters, 6, (2019), 3415–3421.

[16] Pincet, F.; Husson, J. Biophysical Journal, 6, (2005), 4374–4381.

[17] Sitters, G.; Kamsma, D.; Thalhammer, G.; Ritsch-Marte, M.; Peterman, E. J. G.; Wuite, G. J. L. Nature Methods, 1, (2015), 47–50.

[18] Kamsma, D.; Creyghton, R.; Sitters, G.; Peterman, E. J. G.; Wuite, G. J. L. Biophysical Journal, 3, (2016), 501a.

[19] Kamsma, D.; Wuite, G. J. L. Methods in Molecular Biology (Clifton, N.J.) (2018), 341– 351.

[20] Nguyen, A.; Brandt, M.; Muenker, T. M.; Betz, T. Lab on a Chip, 10, (2021), 1929–1947.

[21] Wang, Y. J.; Valotteau, C.; Aimard, A.; Villanueva, L.; Kostrz, D.; Follenfant, M.; Strick, T.; Chames, P.; Rico, F.; Gosse, C.; Limozin, L. Biophysical Journal, 12, (2023), 2518–2530.

[22] Berg-Sørensen, K.; Flyvbjerg, H. Review of Scientific Instruments, 3, (2004), 594–612.

[23] Hyeon, C.; Thirumalai, D. The Journal of Chemical Physics, 5, (2012), 055103.

[24] Chilkoti, A.; Stayton, P. S. Journal of the American Chemical Society, 43, (1995), 10622– 10628.

[25] Dudko, O.; Hummer, G.; Szabo, A. Physical Review Letters, 10, (2006), 108101.

[26] Cossio, P.; Hummer, G.; Szabo, A. Biophysical Journal, 4, (2016), 832–840.

[27] Friddle, R. W.; Noy, A.; De Yoreo, J. J. Proceedings of the National Academy of Sciences, 34, (2012), 13573–13578.

[28] Friddle, R. W.; Podsiadlo, P.; Artyukhin, A. B.; Noy, A. The Journal of Physical Chemistry C, 13, (2008), 4986–4990.

[29] Bullerjahn, J. T.; Hummer, G. Physical Review Research, 3, (2022), 033097.

[30] Petrosyan, R. Journal of Statistical Mechanics: Theory and Experiment, 3, (2020), 033201.

