## Supplementary Figures for "Near equilibrium unbinding of streptavidin/biotin using single molecule acoustic force spectroscopy"

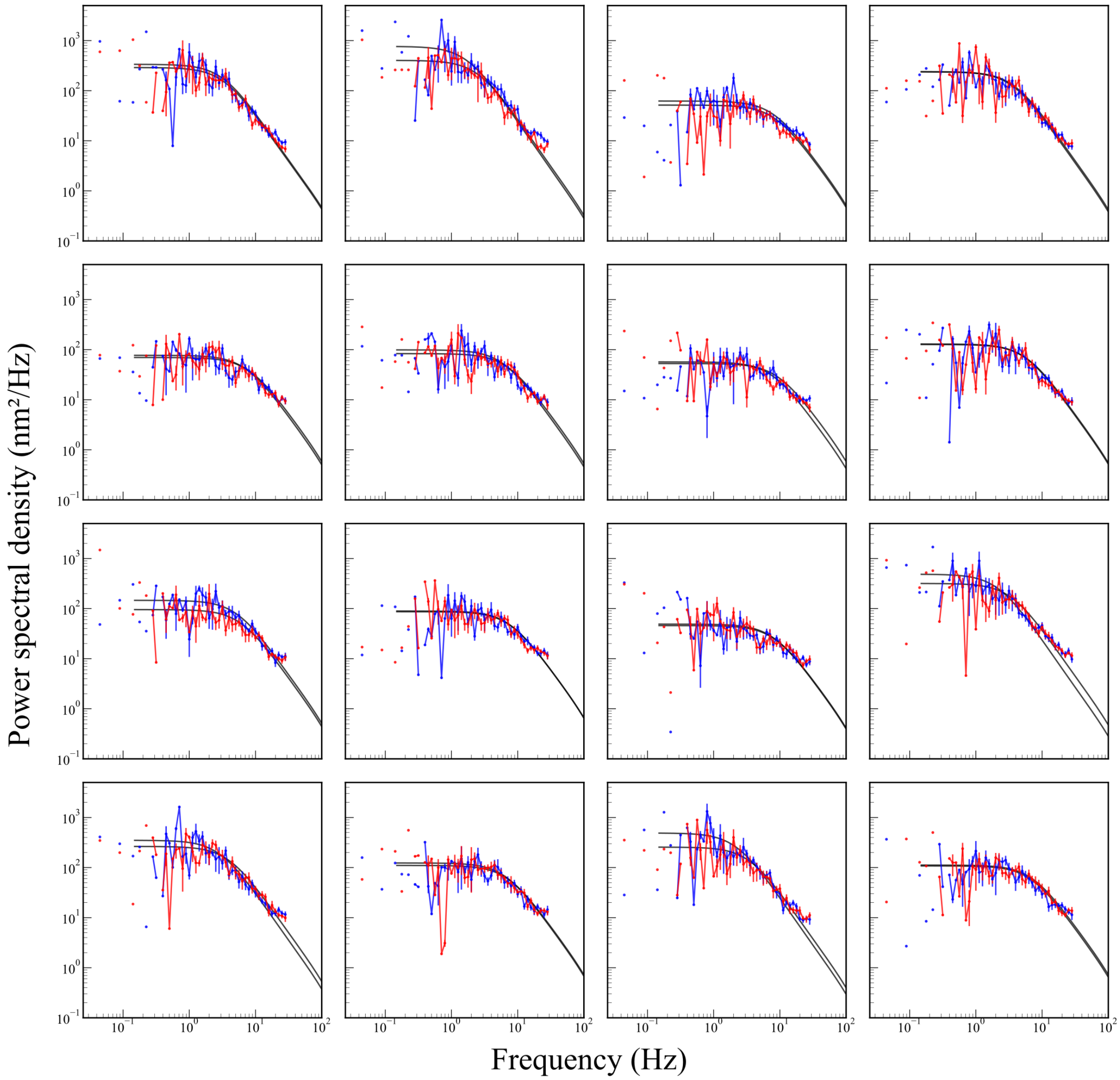


**Figure S1**: **Representative PSD fitting using the OTR strategy.**

Representative Lorentzian fit (black line) to the PSD of both lateral axes (X: blue, Y: red) using the OTR strategy. In the OTR strategy, both the spring constant ($k$) and diffusion constant ($D$) are fitted non-linearly using Equation S1. Each panel represents the fitting for an individual bead.

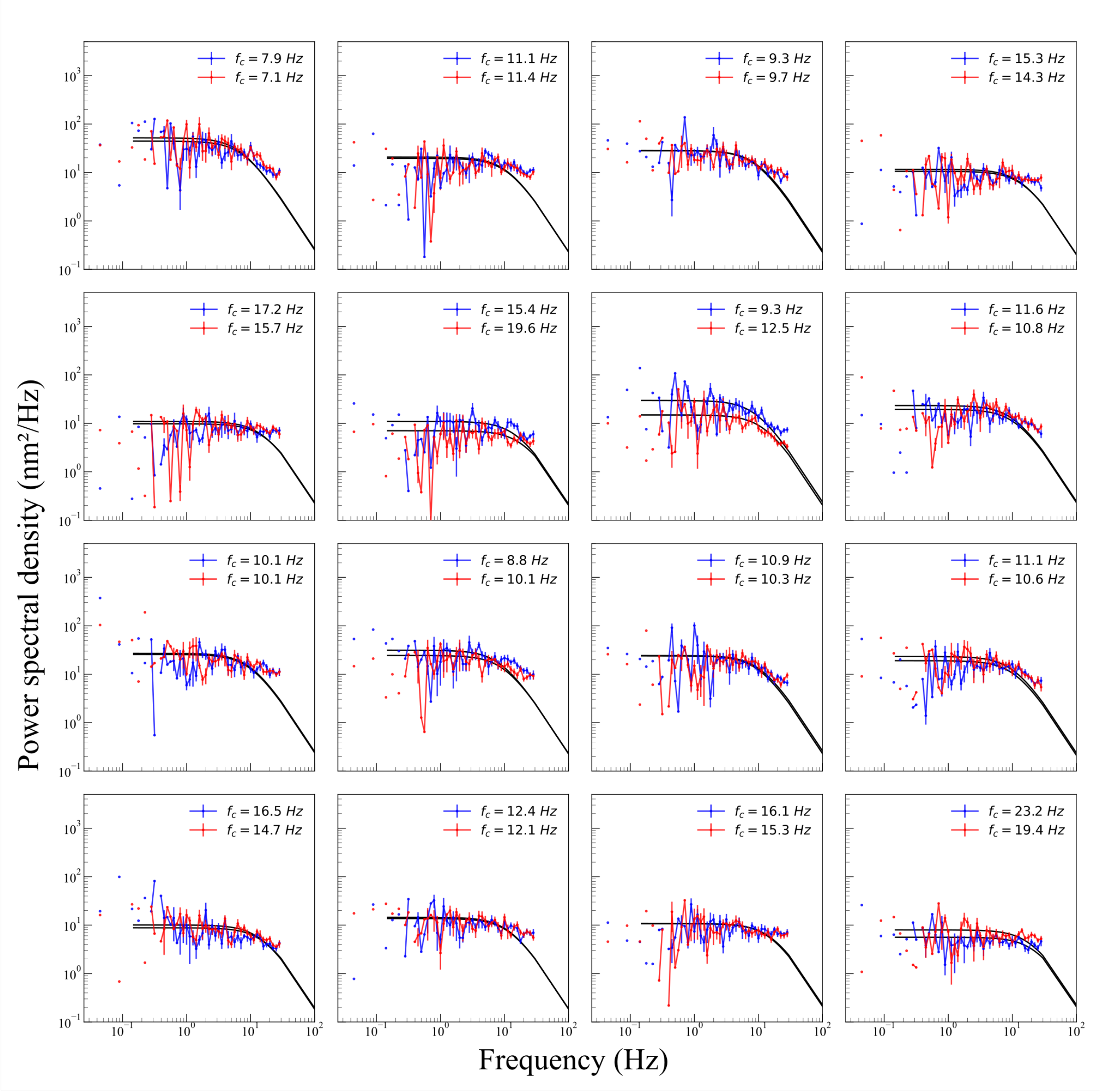


**Figure S2**: **Representative PSD fitting using the OTR-G strategy.**

Representative Lorentzian fit (black line) to the PSD of both lateral axes (X: blue, Y: red). In the OTR-G strategy, an average $D_G$ was determined through non-linear fitting of a rolling OTR strategy until $f_c^{max}$ which was then used in Equation S1, reducing the free fit parameter to only $k$. Each panel represents the fitting for an individual bead.

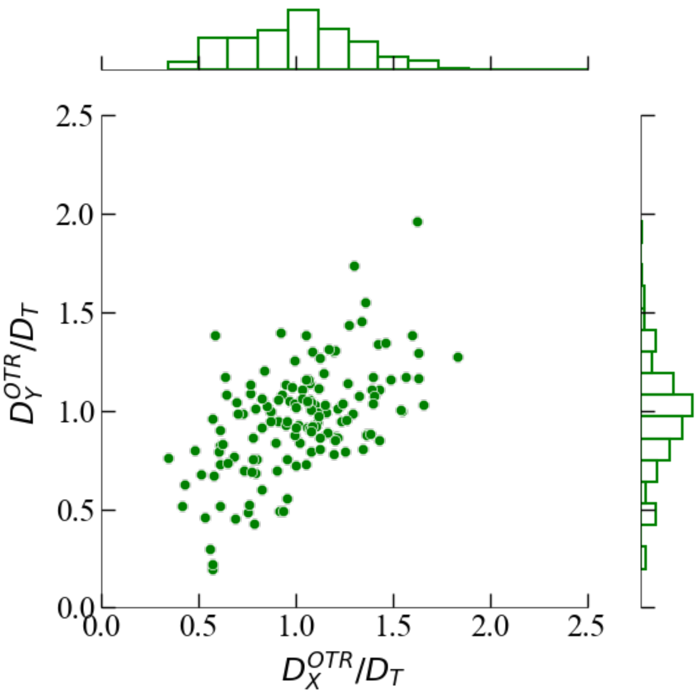


**Figure S3: Ratios of diffusion constants D fitted using the OTR strategy and theoretically expected.**

Scatter plot of the ratios $D^{\mathrm{OTR}}{}_{\mathrm{X}}/D_{\mathrm{T}}$ vs. $D^{\mathrm{OTR}}{}_{\mathrm{Y}}/D_{\mathrm{T}}$, where $D^{\mathrm{OTR}}{}_{\mathrm{X}}$ and $D^{\mathrm{OTR}}{}_{\mathrm{Y}}$ are diffusion constants fitted along the X and Y axes, respectively, using the OTR strategy, and $D_{\mathrm{T}}$ is the theoretical expected diffusion constant. Data shown are from beads for which the PSD was constructed solely using the OTR strategy. A selection criterion for beads to consider was set to a ratio between 0.8 and 1.2.

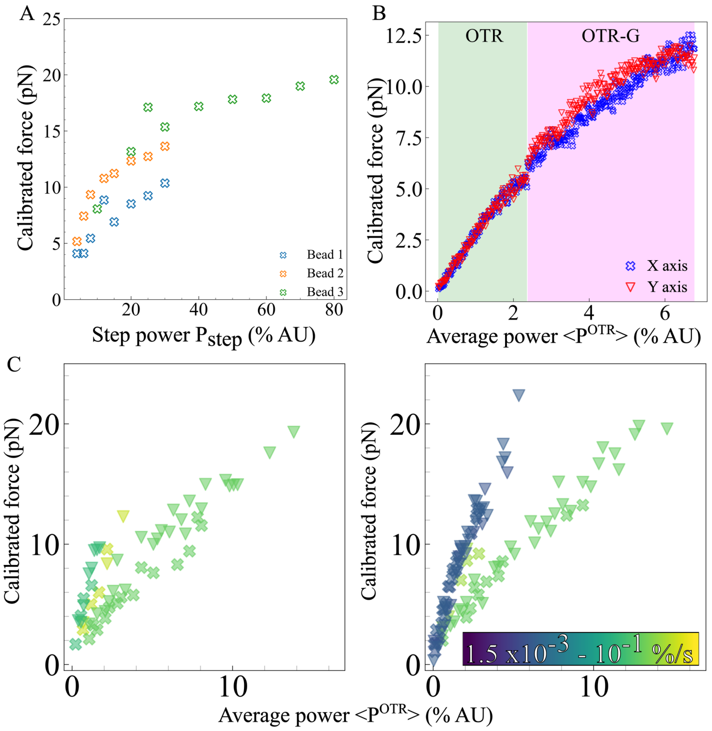


**Figure S4: Non-linear variation in force with power demonstrated by several calibration methods.**

(A) Evolution of the force (calibrated along the X-axis over constant power steps using the traditional method) with the power for three individual beads (different colors) from data recorded at high camera frame rate

(B) Evolution of the force (calibrated along both X (blue) and Y (red) axes using the OTR (green region) and OTR-G (magenta) strategies) with the average power from data recorded at 60 Hz for a single bead.

(C) Evolution of the force (calibrated along both X (crosses) and Y (triangles) axes using the OTR-G strategy) with the power for 3 beads (left) and 4 beads (right) tracked at high camera frame rate (atleast 200 Hz). The color scale indicates the power rate of the linear power ramp.

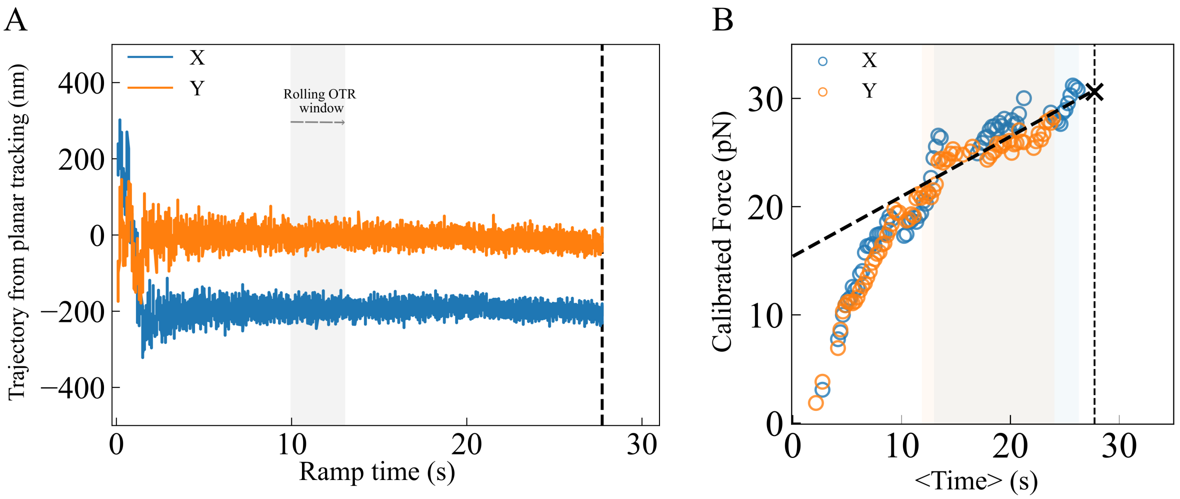


**Figure S5: Strategy to estimate the rupture forces and the loading rate of traces from high pulling rate data.**

(A) Trajectory of a tracked bead resulting from Qitracker implemented on raw videos recorded at 200 Hz for power rates of 1 %/s. The color indicates the lateral axes, and the dotted line represents the bond rupture time, which was determined manually from the raw videos.

(B) Calibrated force versus average time of the OTR window, $<t^{OTR}>$ (i.e. time at which $<P^{OTR}>$ was applied). The nonlinear force versus time is due to nonlinear response of force versus power. The shaded area indicates the regime within which the instantaneous slope was used to estimate the loading rate and the rupture force.

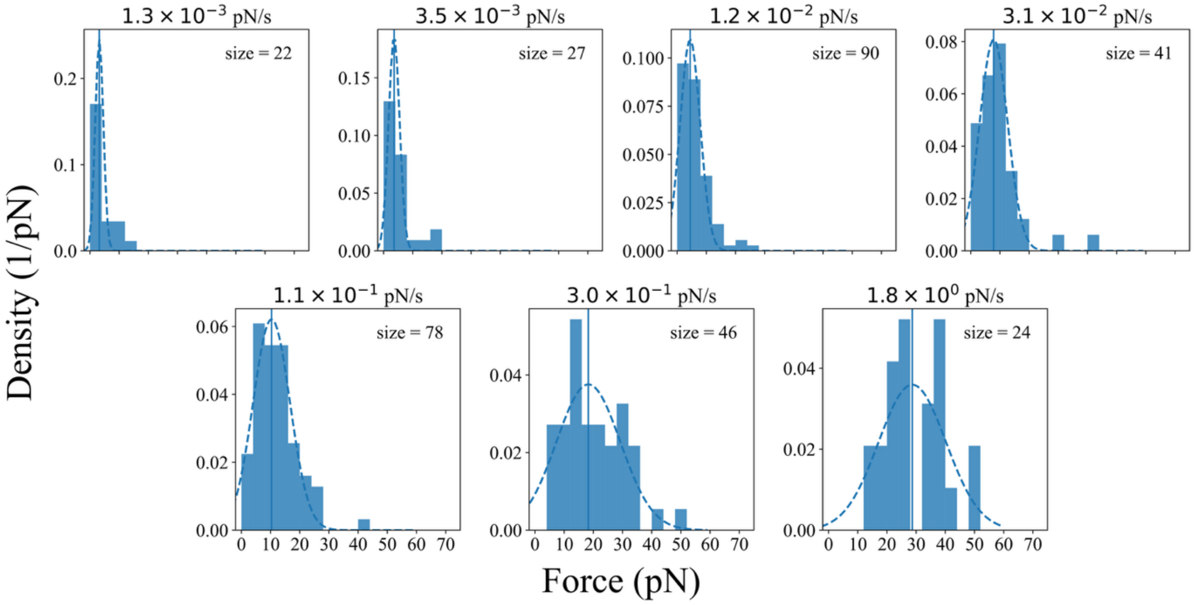


**Figure S6**: Unbinding Force Histograms from AFS experiments

Unbinding force histograms of SA-b from AFS pulling experiments at different loading rates with the corresponding Gaussian fit (dotted line) to extract the most probable unbinding force. Size stands for the number of unbinding events.

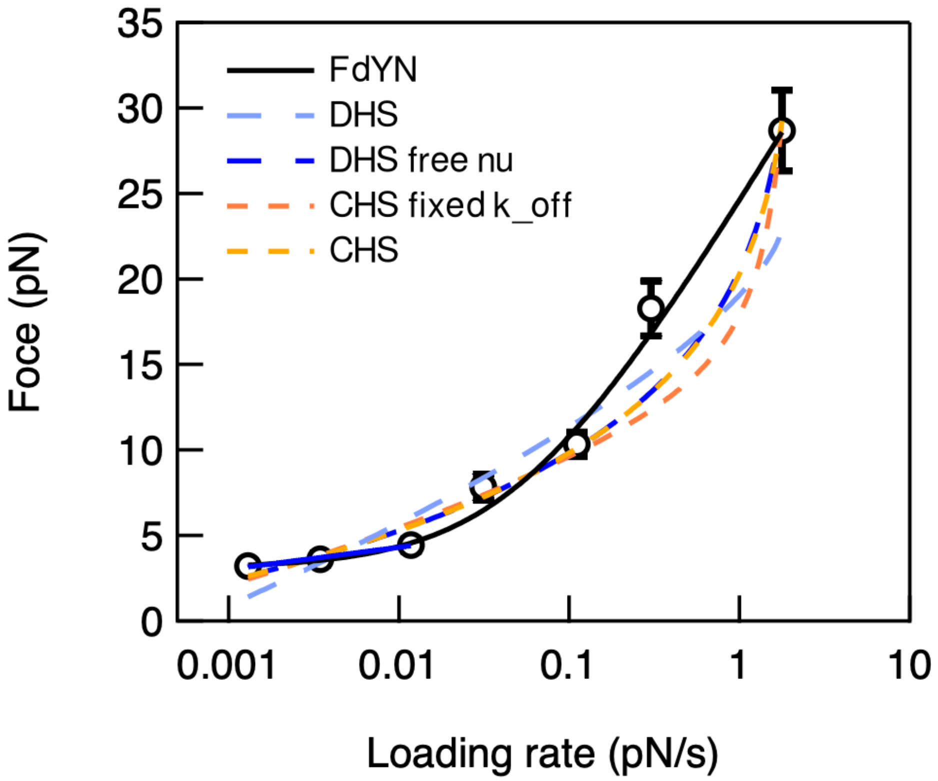


**Figure S7**: Fitting other SMFS theoretical models to AFS data

The DFS data from near equilibrium AFS measurement (open black circles) of the SA-b interaction were fitted with various SMFS models: FNdY (solid black line); CHS (light orange) with a fixed brittleness parameter of 0.01 and a free $k_{off}$; CHS with a free brittleness parameter and a fixed $k_{off}$ of 1.6 x $10^{-5}$ $s^{-1}$ (dark orange); DHS with a fixed landscape shape factor of 2/3 (light blue), DHS with a free landscape shape parameter (dark blue). The obtained fit parameters in each case along with their error estimates are reported in Table S3.

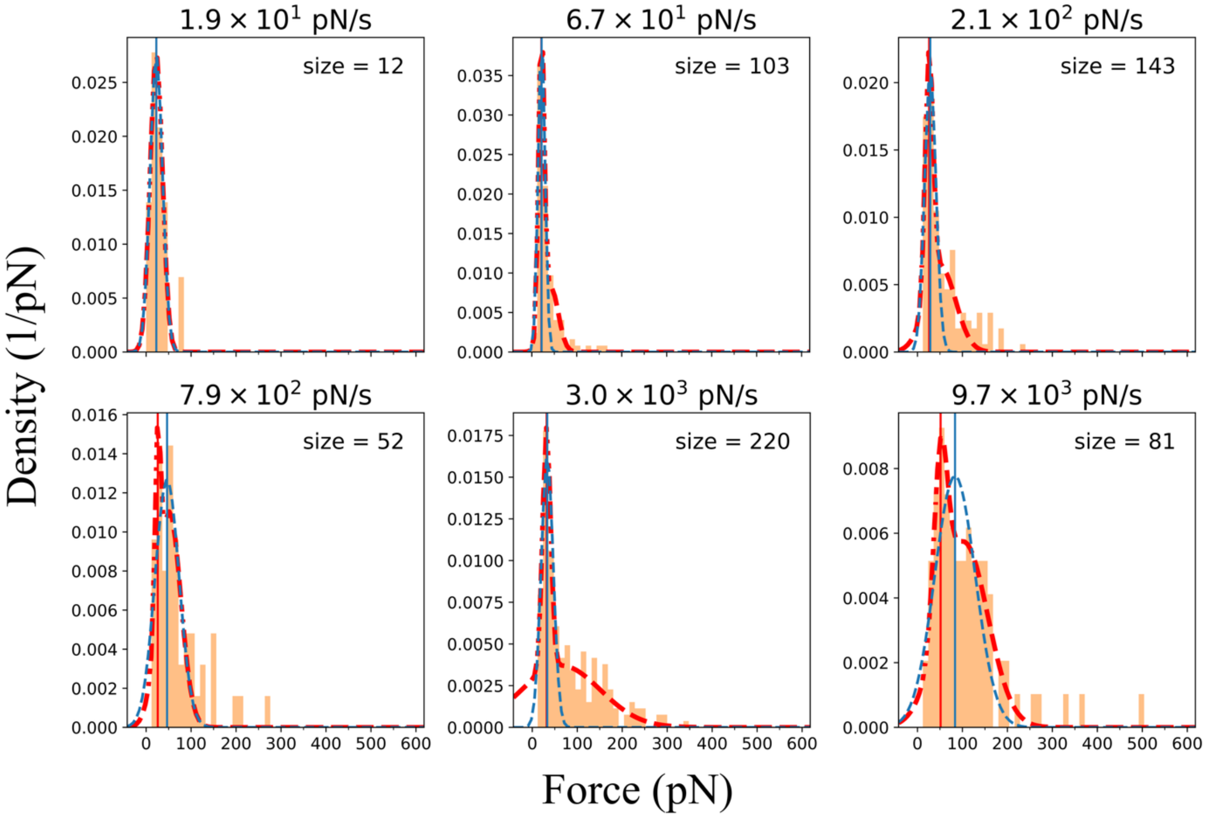


**Figure S8**: Unbinding Force Histograms from AFM experiments.

Unbinding force histograms of SA-b from AFM pulling experiments at different loading rates with the corresponding single (blue dashed line) or bimodal (red dashed line) Gaussian fits to extract the most probable unbinding force. This AFM data complements the AFS near-equilibrium measurements.

**Table S1: Single molecule force spectroscopy measurements of SA-b interaction**

| Techniques | Explored LR range (pN $s^{-1}$) | Model | Number of barriers | $x_\beta$ (nm) | $k_{off}$ ($s^{-1}$) | Reference |
|---|---|---|---|---|---|---|
| Biomembrane force probe | [0.05 – 6 x $10^4$] | Bell Evans | 2 | 0.5<br>0.12 | 0.1-1 | [7] |
| Atomic force microscopy (AFM) | [3 x $10^2$ - 2 x $10^4$] | Bell Evans | 2 | 0.38<br>0.09 | 0.1<br>23 | [10] |
| AFM | [ $10^2$ – $10^5$] | Bell Evans | | 0.46 – 0.073 | 0.72 - 78 | [11] |
| AFM | [ $10^3$ – $10^5$] | Bell Evans | 1, depending on the orientation of SA | 0.41<br>0.23 | 7.7 x $10^{-8}$<br>2.5 x $10^{-8}$ | [15] |
| High speed AFM and Molecular dynamics | [ $10^2$ – $10^{13}$] | Brownian Dynamics | 2 | 0.19<br>0.44 | 1 | [13] |

**Table S2: SA-b dissociation constants using surface plasmon resonance (SPR)**

Equilibrium dissociation constants ($k_{off}$) of SA-b bond were obtained from 5 independent experiments as described in material and methods (SD = standard deviation).

| **Experiment** | **1** | **2** | **3** | **4** | **5** | **Mean ± SD** |
|---|---|---|---|---|---|---|
| $K_{off}$ ($s^{-1}$) | $7.6 \times 10^{-6}$ | $9.6 \times 10^{-6}$ | $1.6 \times 10^{-5}$ | $3.5 \times 10^{-5}$ | $1.3 \times 10^{-5}$ | $1.6 \times 10^{-5} \pm 9.8 \times 10^{-6}$ |

**Table S3: Parameters from other SMFS theoretical models**

| Model | $x_\beta$ (nm) | $k_{off}$ ($s^{-1}$) | Reference |
|---|---|---|---|
| FNdY | 0.54 | 0.003 | [27] |
| CHS, | | | [26] |
| brittleness parameter = 0.01 | 4.8 | $10^{-6}$ | |
| free brittleness parameter | 3 | $1.6 \times 10^{-5}$ | |
| DHS, | | | [25] |
| landscape shape factor =2/3 | 2.1 | $2 \times 10^{-4}$ | |
| free landscape shape factor | 5 | $2.5 \times 10^{-6}$ | |
