## Supplementary material for "Near equilibrium unbinding of streptavidin/biotin using single molecule acoustic force spectroscopy": Materials and Methods

### Reagents and buffers

Double-stranded DNA (1765 base pairs) with an amine group (-NH2) on one end, and a single biotin on the other end were produced by PCR using primers 5'-NH2-GAGTTTTATCGCTTCCATGACG and 5'-Biotin-TTGGTCAGTTCCATCAACATCATAG (Eurogentec) and PhiX174 RF I DNA as template (New England Biolabs). The 1765 base pairs PCR product was purified using Nucleospin columns (Macherey-Nagel) and recovered in 10 mM HEPES, 1 mM EDTA at pH 8.0. The expected contour length of these constructs is 583-600 nm.

The bead coating buffer was composed of 10 mM HEPES, 1 mM EDTA at pH 8.0.

The measurement buffer was composed of 20 mM Tris HCl, 150 mM NaCl at pH 7.8.

### AFS sample preparation, data acquisition & analysis

#### Bead Coating

1 % w/v amino silica beads of 3.1 μm diameter (Spherotech) was washed three times in MilliQ distilled $H_2O$ and incubated in 0.1 % w/v glutaraldehyde for 30 min at room temperature. Beads were then washed three times in coating buffer and then incubated in a diluted solution of biotinylated DNA (1.7 pM) in coating buffer for at least 2 hours. The DNA-coated beads were washed three times and stored at 4°C in the measurement buffer for up to 1 week.

#### AFS chamber functionalization

Before each experiment, the AFS chamber was cleaned three times with 200 μL of 1 M NaOH (Sigma-Aldrich) for 2 minutes and then rinsed with 1 mL of MilliQ. The chamber surface was activated by three successive incubations with 200 μL of 1M HCl (Sigma-Aldrich) for 2 minutes, and then rinsed with 1 mL of $H_2O$. Bare silica beads of diameter 1.6 µm (Spherotech) were washed three times in MilliQ and 40 µL of a 0.025 % w/v solution was injected in the chamber and incubated for 20 minutes at room temperature (RT). These beads adhered to the surface of the chamber and were used as a reference to correct for thermal drift in the chamber. The chamber was then filled with 40 µL of streptavidin (Sigma-Aldrich) at 20 µg/mL in 100 mM of $NaHCO_3$ buffer and incubated for 60 minutes at RT. After rinsing with measurement buffer, the chamber was passivated by incubation with 40 µL of 0.2 % w/v bovine serum albumin (BSA, New England Biotech lab) in measurement buffer for 60 minutes at RT to minimise non-specific interactions. DNA coated beads were also passivated with 0.2 % w/v BSA for 60 minutes at RT. The passivated DNA coated beads were injected in the chamber at 0.02 % w/v for optimal bead tracking.

**AFS data acquisition**

The AFS generator (Lumicks, generation 3) was controlled by the manufacturer's LABVIEW software (Lumicks AFS Tracking v1.4.0). The AFS chamber (B chamber, pulling frequency ~7.5 MHz, Lumicks) was mounted on a motorized stage (MLS203, Thorlabs) and illuminated by a fibre-coupled LED (M660F1, Thorlabs). The beads were imaged with a 20x air objective (Uplan F, Olympus) using a USB camera (UI1324, IDS) at a frame rate ($f_s$) of 60 Hz and exposure time ($\tau_e$) of 2.3 ms.

A few reference beads and a few tens of DNA-tethered beads were selected (non-moving ROI of 50 pixels). A look-up-table was predefined ranging from 0 to 10 µm in steps of 0.1 µm, allowing 3D tracking. During experiments, the bonds were loaded using a linear power ramp at a defined rate (dP/dt in units of % per second), with maximum power typically up to 7% (AU), were applied for durations ranging from 600 seconds to 12 hours. The individual bead trajectories were recorded as a TDMS file (i.e. LABVIEW's native format) along with power amplitude, chip drive frequency and temperature as a corresponding time series.

For high pulling rates (i.e. power rate of 1 %/s), as the power increases rapidly and results in short bond survival times, a CMOS camera (acA720-520 µm, Basler) with a higher frame rate (200 Hz) was used and videos of the entire field of view were recorded. 2D bead trajectories were tracked offline using a code adapted from Qitracker (Fig S5)[1].

### AFS data analysis:

#### Correction for Z coordinates

Since an air objective was used to measure the apparent height of beads in liquid, a correction of the Z position based on the lookup table is required. To determine the real height of the beads, the difference in refractive index (*n*) between water and air has to be accounted, through a multiplicative height correction factor $n_{water}/n_{glass} = 1.33$[2].

#### Traces selection and analysis

Bead trajectories showing a single tether signature were manually selected based on the increase in height in the Z-direction during power application and the in-plane circular motion at zero power[3]. We analysed the selected beads by adapting to linear power ramp experiments the Python-based data analysis pipeline previously developed for power clamp experiments[4].

Briefly, the traces were corrected for drift using averaged reference beads traces. The anchor point in XY plane was determined by taking the average position over a 3 minutes time window at zero force at the beginning of the trace, while the anchor point in Z was defined as the minimum Z position of the trace. The bond rupture power $P_{r,}$ at which the bead detached was identified as the loss of the tracking signal, determined manually through the GUI to avoid mislabelling tracking errors as rupture events.

For the traces acquired at a higher frame rate (200 Hz) and tracked offline, the rupture frame number for each bead was also determined manually from the recorded videos since the offline tracking was shown to be more prone to tracking errors.

Determination of the loading rate and rupture force at low power ramp

Once the rupture power has been identified, a time window with of $N_{PSD}$ = 1200 data points (i.e 20 seconds for $f_s$ = 60 Hz) was manually defined on the linear power ramp close to the unbinding point, while avoiding potential spikes or tracking artefacts in the traces. A one-sided Power Spectral Density (PSD) of the horizontal fluctuations of the bead was constructed along each axis using the periodogram function of the Python SciPy[5] module. For graphical display, the PSD was binned into 101 equal logarithmically spaced bins on the frequency axis (Fig 1C & D, Fig S2 & S3).

If the corner frequency $f_c$ which separates the elastic and the diffusive regime in the PSD was visually observable and less than $f_s/6$ with $f_s$ the camera sampling rate. The PSD was fitted with the thermal spectral density of a diffusive bead trapped in a harmonic potential of stiffness $k$ according to Eq S1[6] (which also accounts for both motion blurring and aliasing effects through the summation of the sine components[7]).

$$PSD(f) = \sum_{n=0}^{1} \frac{4(k_B T)^2}{Dk^2} \frac{1}{1+\left((f+nf_s)/f_c\right)^2} \frac{sin^2\left(\pi\tau_e(f+nf_s)\right)}{\left(\pi\tau_e(f+nf_s)\right)^2} \qquad \text{(Eq S1)}$$

with $D$ the bead diffusion coefficient, $k$ the stiffness of the inverted pendulum along a horizontal axis, fc the corner frequency that delineates the elastic and diffusive parts of the PSD, $k_B T$ the thermal energy, $f_s$ the camera sampling rate and $\tau_e$ the exposure time. The fitted diffusion coefficient $D^{OTR}$ was compared to the theoretical estimate $D_T$ taking into account the hydrodynamic interaction between the chamber wall and the bead of radius $R$ at height $Z$[8]:

$$D_T^{faxen} = \frac{k_B T}{6\pi\eta R}\left[1 - \frac{9}{16}\left(\frac{R}{Z+R}\right) + \frac{1}{8}\left(\frac{R}{Z+R}\right)^3\right]^{-1} \quad \text{(Eq S2 )}$$

The ratio $D^{OTR}/D_T$ was used to assess the goodness of the PSD fit along each axis for a single tethered bead (Fig S3). The selection threshold for this ratio was set within [0.8,1.2].

If the corner frequency $f_c$ was more than $f_s/6$, the OTR-G strategy was implemented. As $D$ can be assumed to be an intrinsic bead property, an average $D_G$ along each axis was calculated from the PSD fit over rolling windows every 120 points (10 % $N_{PSD}$) at the beginning of the power ramp, for which $f_c$ is less than $f_s/6$ and $D_{fit}/D^{faxen}_T$ met the selection threshold. $D_G$ was substituted back into the Eq S1, thus reducing the fit of the pre-rupture PSD to a single unknown, k.

From the fitted $k_x$ and $k_y$ for the two lateral axes, an average vertical pulling force $<F^{OTR}>$ experienced by the bead was calculated as

$$< F^{OTR} >= \frac{(k_x + k_y)}{2}(L + R) \quad \text{(Eq S3)}$$

with R the radius of the bead and L the DNA tether length calculated from the tracked data[4] (SFig S1). This force is attributed to the average power $<P^{OTR}>$ exerted on the bead during the time window used to construct the PSD.

The unbinding force $F_u$ and its corresponding instantaneous loading rate $r_f$ are deduced from $<F^{OTR}>$ and $P_r$ which the bond ruptured using the following equations

$$F_u = \frac{< F^{OTR} >}{< P^{OTR} >} \times P_r \quad \text{(Eq S4)}$$

$$r_f = \frac{< F^{OTR} >}{< P^{OTR} >} \times \frac{dP}{dt}$$

Determination of the loading rate and rupture force at high power ramp

At high powers, the force-power relationships were non-linear. Hence, for traces acquired at high pulling rate (1%/s) with high sampling rates (200 Hz) and tracked offline, the PSDs were built using only $N_{PSD}$ = 600 data points (i.e. a window of 3 seconds). To exhaustively describe the non-linear force-power relation of each individual bead, PSDs in X and Y axes were built using a rolling window every 60 points (10 % $N_{PSD}$ ) and subsequently fitted using the OTR-G strategy. This resulted in a force-power plot for each bead along each axis (Fig S5). All the force values showing negative correlation with power or exhibiting random jumps typically due to incorrect PSD fitting or tracking errors were removed manually.

A linear fit of the last 50% of the force-power plot was used to extrapolate the force at the unbinding point and to estimate the instantaneous loading rate (slope of the force versus time fit) (Fig S5). Additionally, beads with $P_r$ exceeding 30% where the force might not be increasing linearly were excluded from the analysis.

Code Availability

All the data analysis routines presented in this work were implemented using custom-written modular Python code, and a simple graphical user interface was also built for exploratory analysis. The code is available on the GitHub repository (source code: https://github.com/DyNaMo-INSERM/PyAFSpectra).

### AFM sample preparation, data acquisition & analysis

Streptavidin-coated agarose beads (Sigma-Aldrich) were immobilized on the sample surface by embedding them in a thin 4% agarose layer. Biotin was covalently attached to the AFM cantilever (MLCT-bio, Bruker) through a PEG linker (contour length~10 nm). Briefly, AFM probes were rinsed with water and acetone for 10 min, dried with nitrogen, cleaned for 15 min by UV-ozone and incubated for 10 min in 5% APDMES in ethanol (abcr GbmH). The AFM probes were rinsed in ethanol, dried over a gentle flow of nitrogen, baked at 80°C for 30 min and incubated overnight in 100mM sodium borate buffer at pH 8. The amino-silanised probes were immersed in a solution of 1 mg/ml NHS-PEG- biotin (3kDa, Sigma-Aldrich) for 1 h. The probes were then rinsed with phosphate buffer saline (PBS) at pH 7.4 and stored at 4 °C until use.

AFM measurements were performed using a Nanowizard 4 (JPK, Germany). The experiments were conducted at room temperature in the measurement buffer. During force curve acquisition, the cantilever was approach at 2 µm/s and retracted at velocities varying from 0.02 µm/s to 2 µm/s with a contact time of 100 ms. The optical lever sensitivity was calibrated by acquiring a force curve on a hard surface at the end of the experiment and the spring constant was determined using the thermal spectrum in liquid[9].

The resulting force curves were analyzed using the software provided by the manufacturer (JPK Data Processing). The rupture force was defined as the force of the last adhesion peak, and the loading rate was defined as the slope of the last portion of the peak on force curves plotted versus time. The adhesion peaks with a contour length lower than 10 nm or larger than 50 nm were discarded.

### Surface Plasmon Resonance data acquisition and analyses

Binding experiments and kinetic analysis were performed at 22 °C using a Biacore T200 instrument (Cytiva) using measurement buffer as the running buffer. Streptavidin was coupled on two consecutive flow cells (experimental and control) of a CM3 sensor chip (Cytiva) according to the manufacturer's instructions. Briefly, the matrix was chemically activated by the injection of a 1: 1 mixture of N-hydroxysuccinimide and 1-ethyl-3-(3-dimethylaminopropyl)carbodiimide. Streptavidin (SA) (100 µg/ml) was then injected at pH 4.5. Non-occupied activated sites were blocked using a 1 M ethanolamine injection followed by a pulse of 2M NaCl to remove non-covalently bound streptavidin. Under these conditions, about 700 RU of streptavidin were coupled on each of the experimental and control flow cells.

The stability of the background signal was monitored by automatically measuring the signal differences between experimental and control flow cells. Once stabilized, biotinylated DNA (12 nM) was injected (5 µl/min) over the experimental flow cell (70-190 RU) and five consecutive washing steps with running buffer (8s at 40 µl/min) were performed. DNA dissociation data were collected over at least 7200 seconds. In all cases the global dissociation was above 5% of captured DNA values. SPR traces obtained during DNA dissociation were deduced from background signal to calculate dissociation rate constants ($k_{off}$). The $k_{off}$ was determined using the Biaevaluation 3.1 software using a first-order exponential decay (Table S2). All standard errors (SE) of the theoretical fits given by the software during $k_{off}$ calculation were lower than 1% of the calculated $k_{off}$ value.

### REFERENCES


[1]. van Loenhout, M. T. J.; Kerssemakers, J. W. J.; De Vlaminck, I.; Dekker, C. Biophysical Journal , 10, (2012), 2362–2371.

[2]. Hell, S.; Reiner, G.; Cremer, C.; STELzer, E. H. K. Journal of Microscopy , 3, (1993), 391–405.

[3]. Nelson, P. C.; Zurla, C.; Brogioli, D.; Beausang, J. F.; Finzi, L.; Dunlap, D. The Journal of Physical Chemistry. B , 34, (2006), 17260–17267.

[4]. Wang, Y. J.; Valotteau, C.; Aimard, A.; Villanueva, L.; Kostrz, D.; Follenfant, M.; Strick, T.; Chames, P.; Rico, F.; Gosse, C.; Limozin, L. Biophysical Journal , 12, (2023), 2518–2530.

[5]. Virtanen, P.; Gommers, R.; Oliphant, T. E.; Haberland, M.; Reddy, T.; Cournapeau, D.; Burovski, E.; Peterson, P.; Weckesser, W.; Bright, J.; van der Walt, S. J.; Brett, M.; Wilson, J.; Millman, K. J.; Mayorov, N.; Nelson, A. R. J.; Jones, E.; Kern, R.; Larson, E.; Carey, C. J.; Polat, İ.; Feng, Y.; Moore, E. W.; VanderPlas, J.; Laxalde, D.; Perktold, J.; Cimrman, R.; Henriksen, I.; Quintero, E. A.; Harris, C. R.; Archibald, A. M.; Ribeiro, A. H.; Pedregosa, F.; van Mulbregt, P. Nature Methods , 3, (2020), 261–272.

[6]. Berg-Sørensen, K.; Flyvbjerg, H. Review of Scientific Instruments , 3, (2004), 594–612.

[7]. Wong, W. P.; Halvorsen, K. Optics Express , 25, (2006), 12517–12531.

[8]. Leach, J.; Mushfique, H.; Keen, S.; Di Leonardo, R.; Ruocco, G.; Cooper, J. M.; Padgett, M. J. Physical Review E , 2, (2009), 026301.

[9]. Sumbul, F.; Rico, F. Single-Molecule Force Spectroscopy: Experiments, Analysis, and Simulations. In *Atomic Force Microscopy: Methods and Protocols*; Santos, N. C., Carvalho, F. A., Eds.; Springer: New York, NY, 2019; pp 163–189.
